## Supplementary information for "Phylogenomics reveals the evolutionary origins of lichenization in chlorophyte algae"

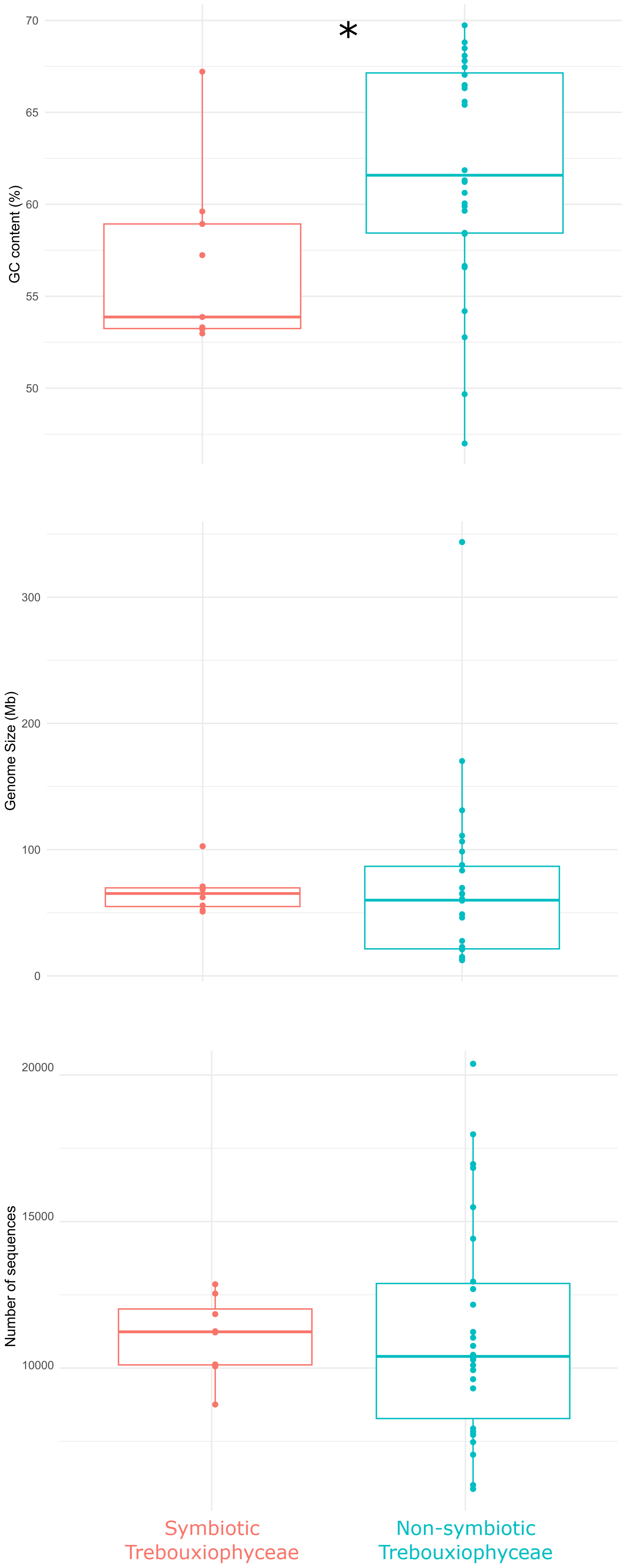


**Supplementary Fig. 1: Genomic comparisons including the GC content, the genome size, and the number of sequences between symbiotic Trebouxiophyceae (pink, n=13) and non-symbiotic Trebouxiophyceae (blue, n=23).** Statistical comparisons have been performed using a two-sided Wilcoxon test and p-value threshold of 0.05. The first (minimum) and third (maximum) quartiles and the median are presented in the boxplots

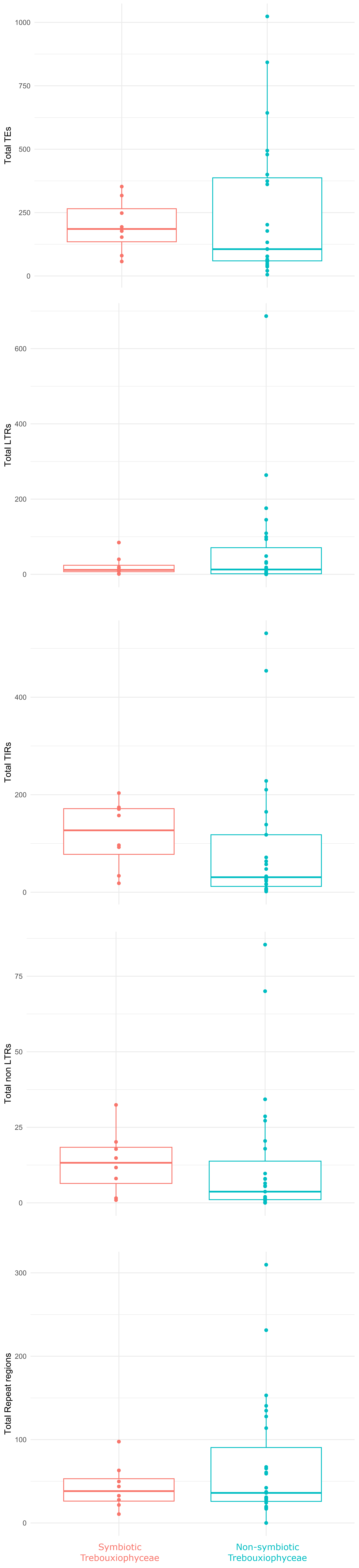


**Supplementary Fig. 2: Comparison of transposable elements percentage between symbiotic Trebouxiophyceae (pink, n=13) and non-symbiotic Trebouxiophyceae (blue, n=23).** Statistical comparisons have been performed using a two-sided Wilcoxon test and p-value threshold of 0.05. using a two-sided Wilcoxon test and p-value threshold of 0.05. The first (minimum) and third (maximum) quartiles and the median are presented in the boxplots

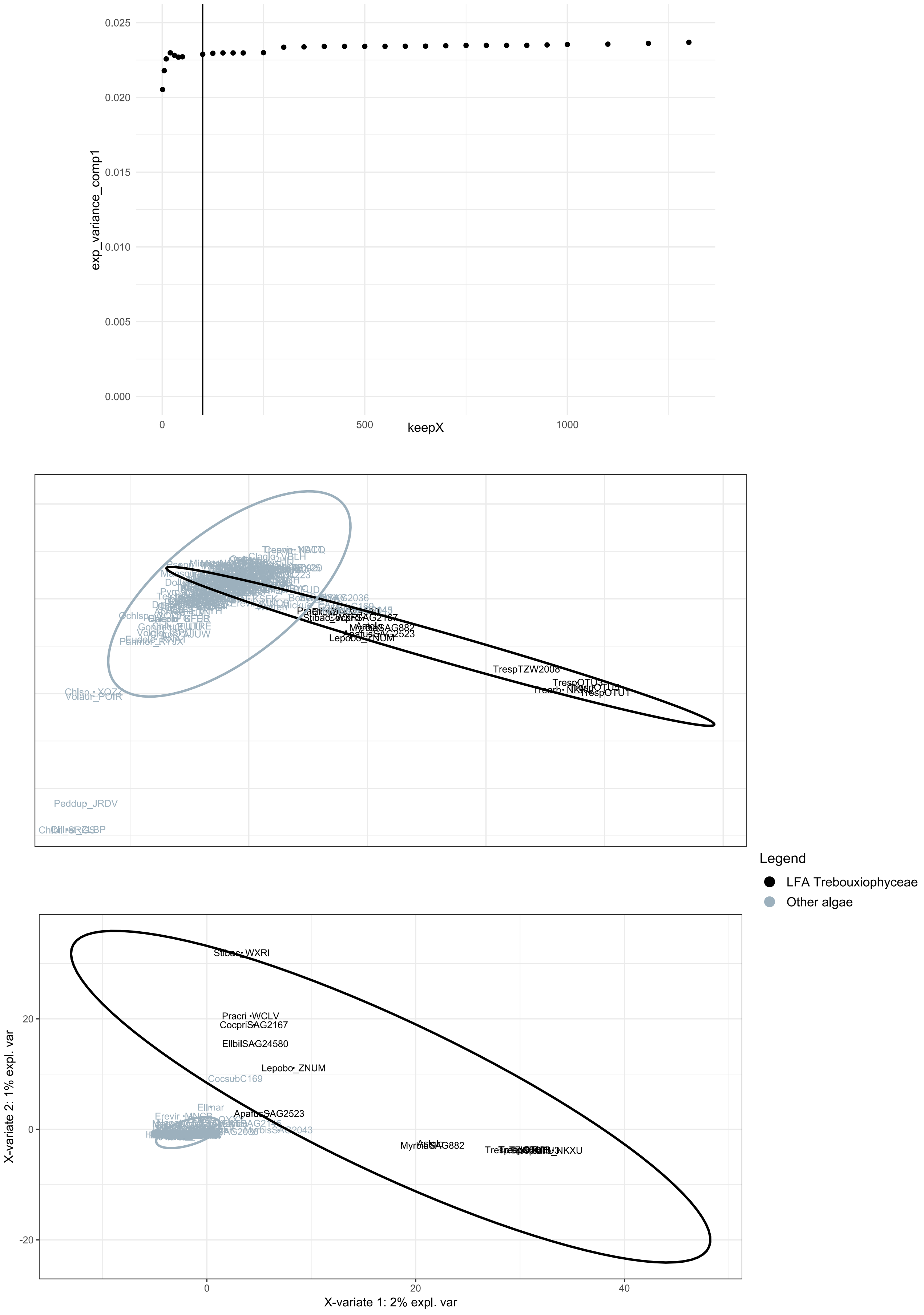


**Supplementary Fig. 3: PLS-DA results** (**a**) Explained variance for the first component depending on the number of explicative orthogroups. The threshold was set to 100 orthogroups. (**b**) Individual plot for the PLS-DA by taking all the orthogroups into account ( **c**) Individual plot for the sparse PLS-DA based on the 100 best orthogroups.

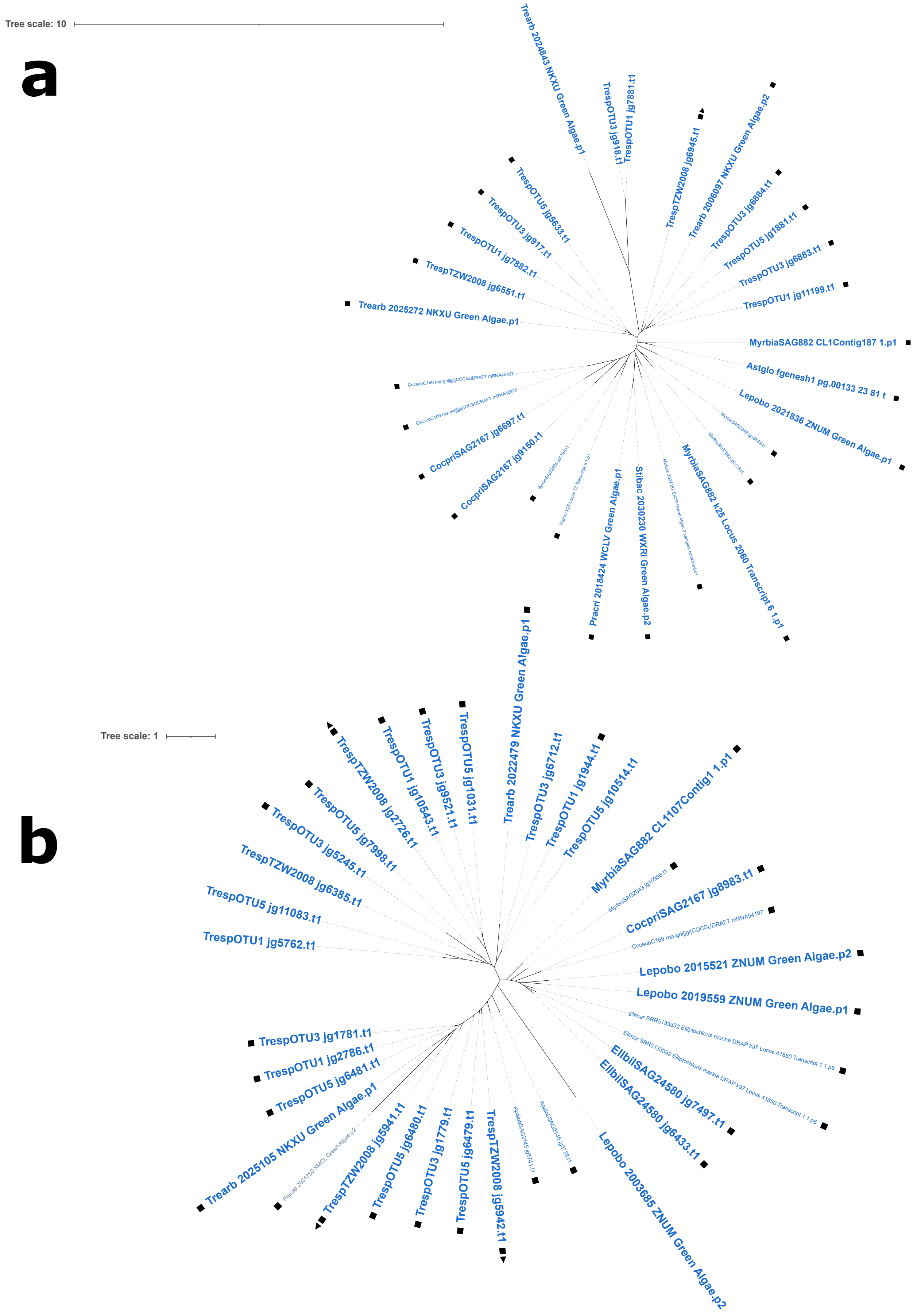


**Supplementary Fig. 4: Phylogenetic trees of the putative horizontal gene transfer candidates using the genomes and transcriptomes database described in Table S1.** (**a**) Unrooted phylogenetic tree of N0.HOG00012965 (Glycoside hydrolase 8). (**b**) Unrooted phylogenetic tree of N0.HOG00012501 (glutathione S-transferase). Trebouxiophyceae are shown in blue, the other Chlorophytes (sensu lato) are shown in grey. Lichen forming species are shown in bold. Black squares materialize the initial orthogroup composition given by OrthoFinder and the black triangle shows the differentially expressed gene during the symbiosis between TrespTZW2008 and *Usnea hakonensis*

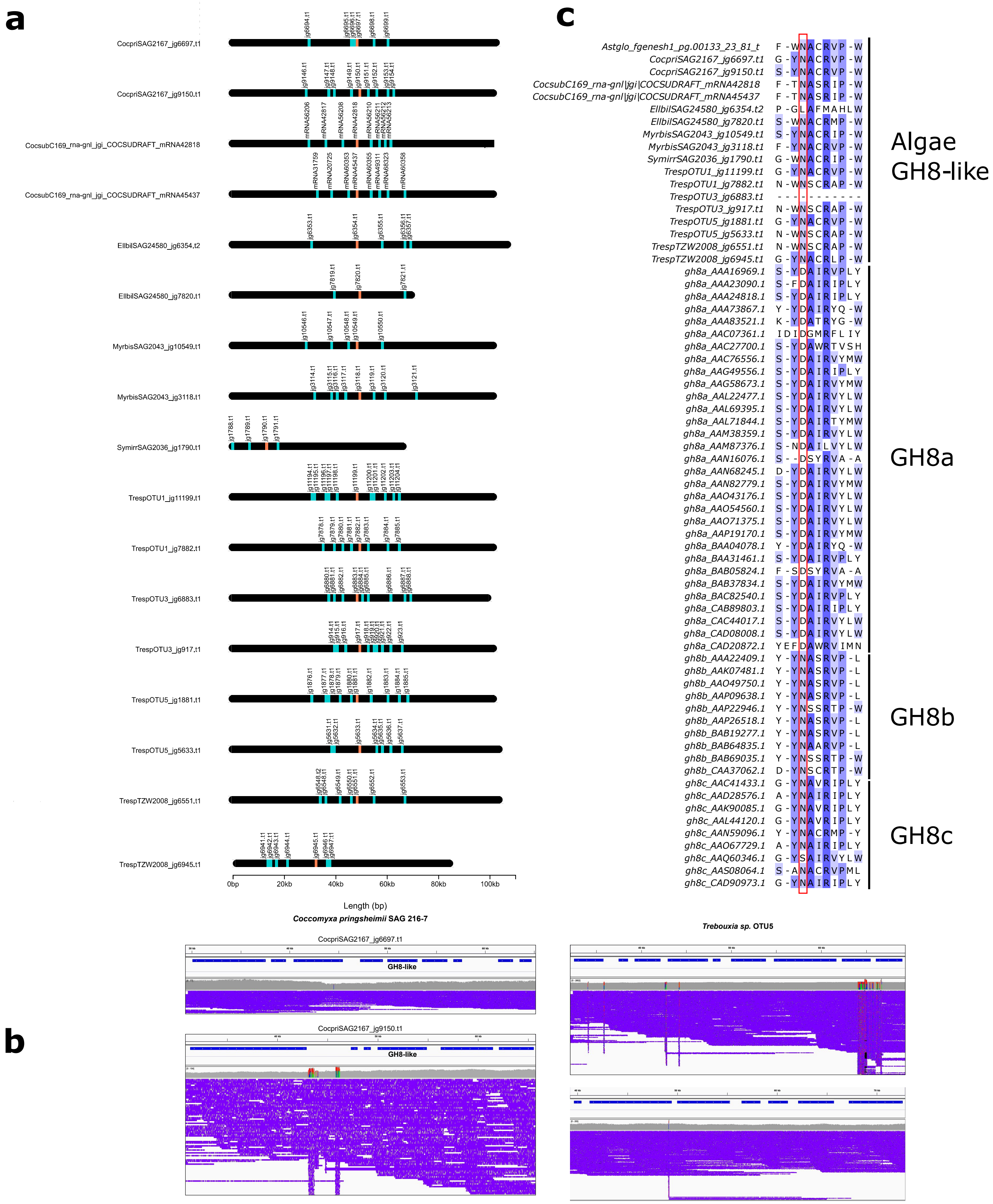


**Supplementary Fig. 5: Horizontal gene transfer demonstration for the GH8 enzyme.** (**a**) Genomic context on 100kb regions centered on algae GH8 (coral bars). Other genes in these regions are marked with cyan bars. ( **b**) Reads mapping on a 100kb region of *Trebouxia sp* (OTU5) and *Coccomyxa pringsheimii* . ( **c**) Alignment of GH8 subfamilies catalytic sites (highlighted by the red square). Bacterial sequences are from Adachi et al, 2004.

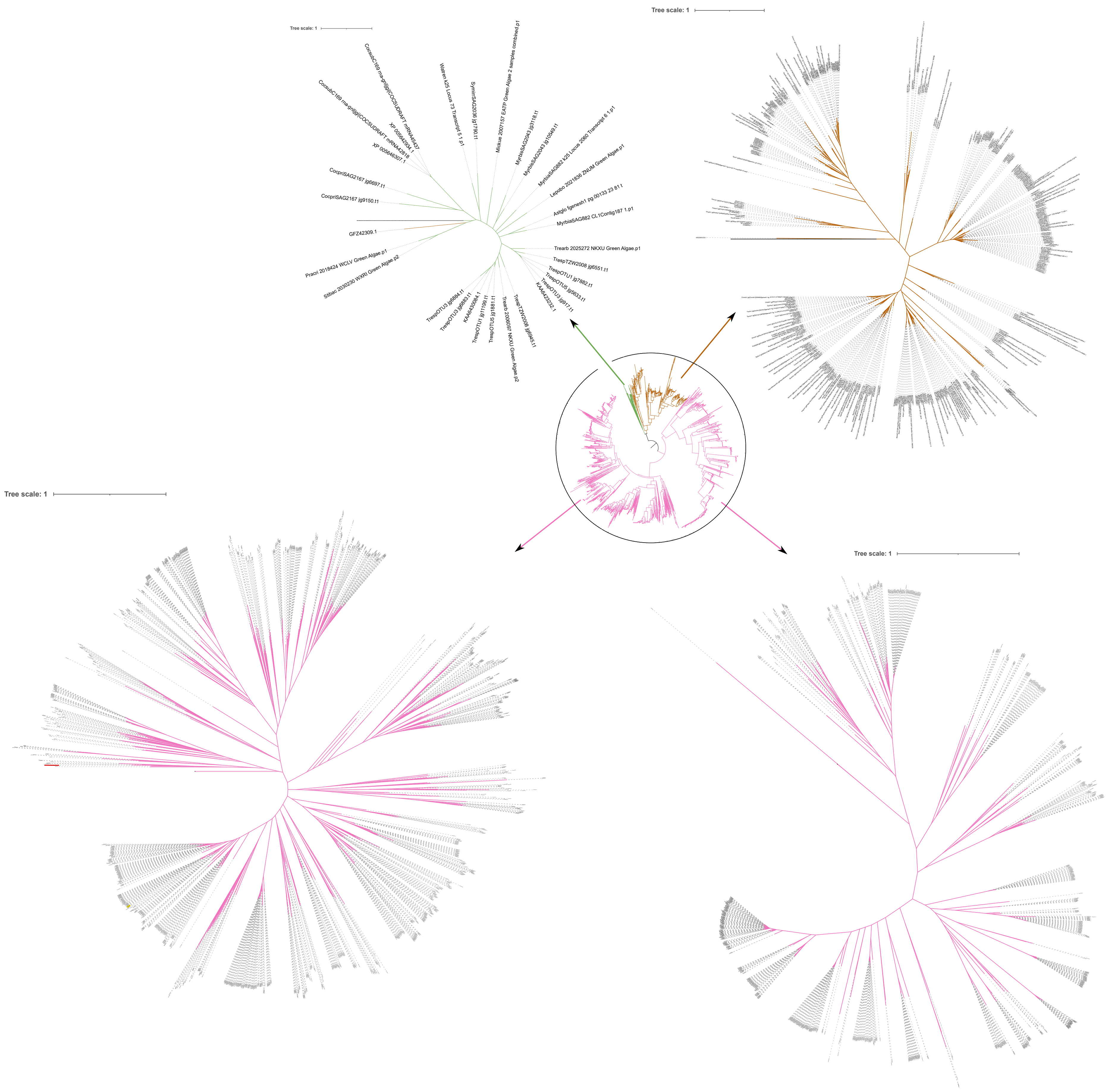

**Supplementary Fig. 6: Maximum Likelihood tree of the GH8 proteins.** Branches are colored as follow: bacteria in pink, fungi in brown and chlorophytes algae in green.

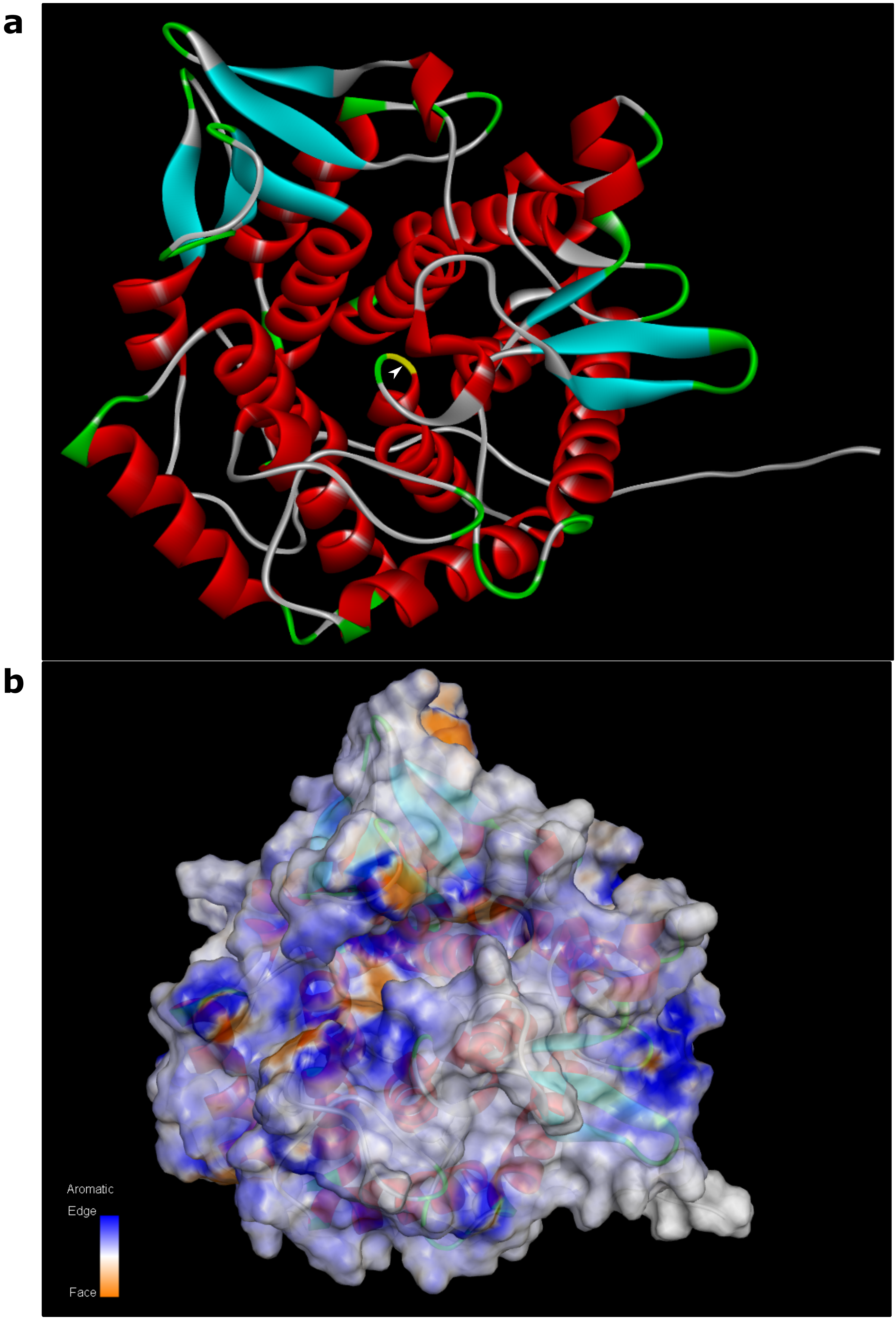


**Supplementary Fig. 7: AlphaFold prediction of the *A. glomerata* GH8.** (**a**) 3D structure of the GH8 with the active catalytic site indicated with a white arrow. (**b**) Aromatic structure of the GH8 with the putative binding pouch containing the active catalytic site.

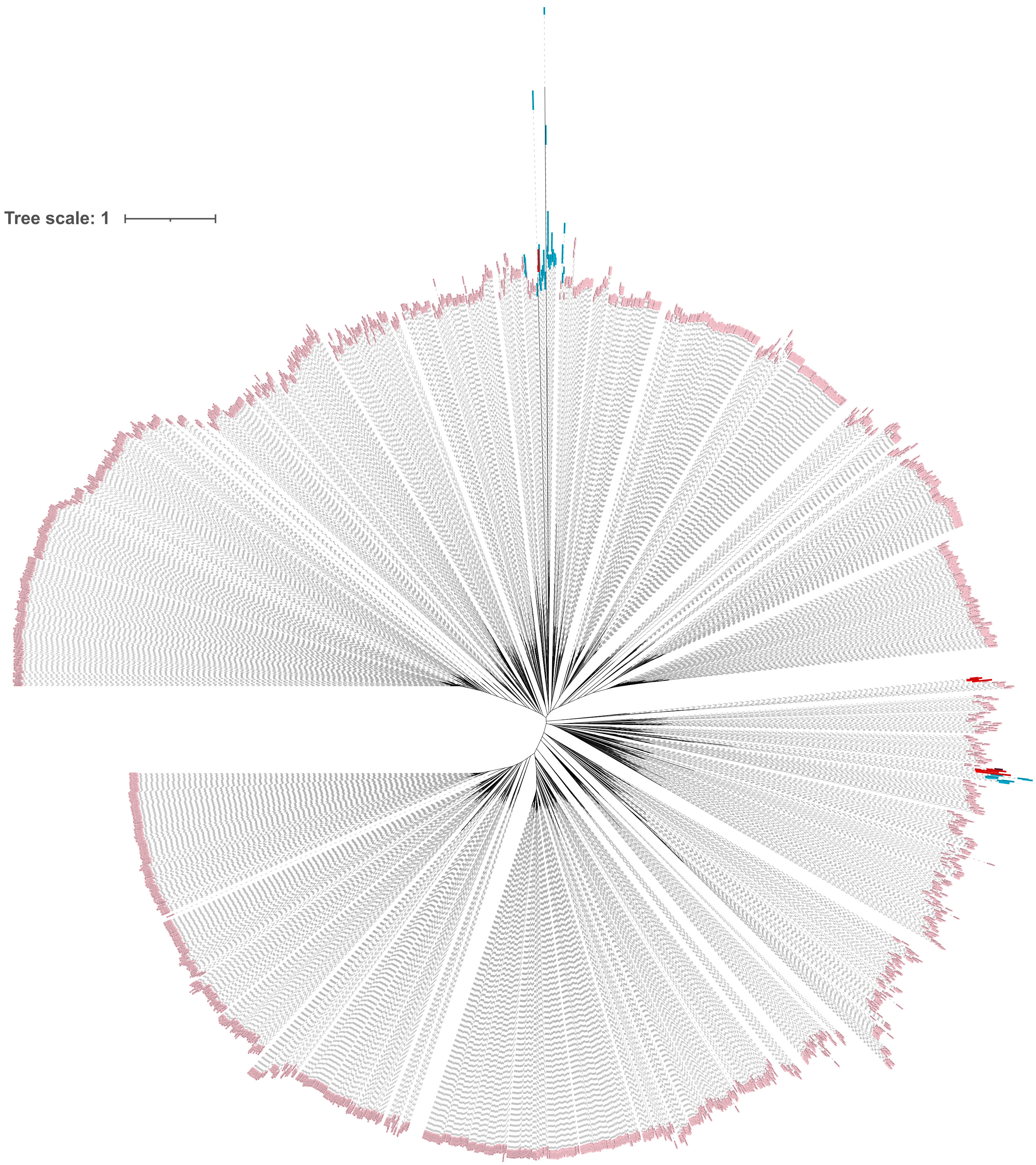


**Supplementary Fig. 8: Maximum Likelihood tree of the glutathione S -transferase proteins.** Labels are colored as follow: bacteria in pink, fungi in red and chlorophytes algae in blue.

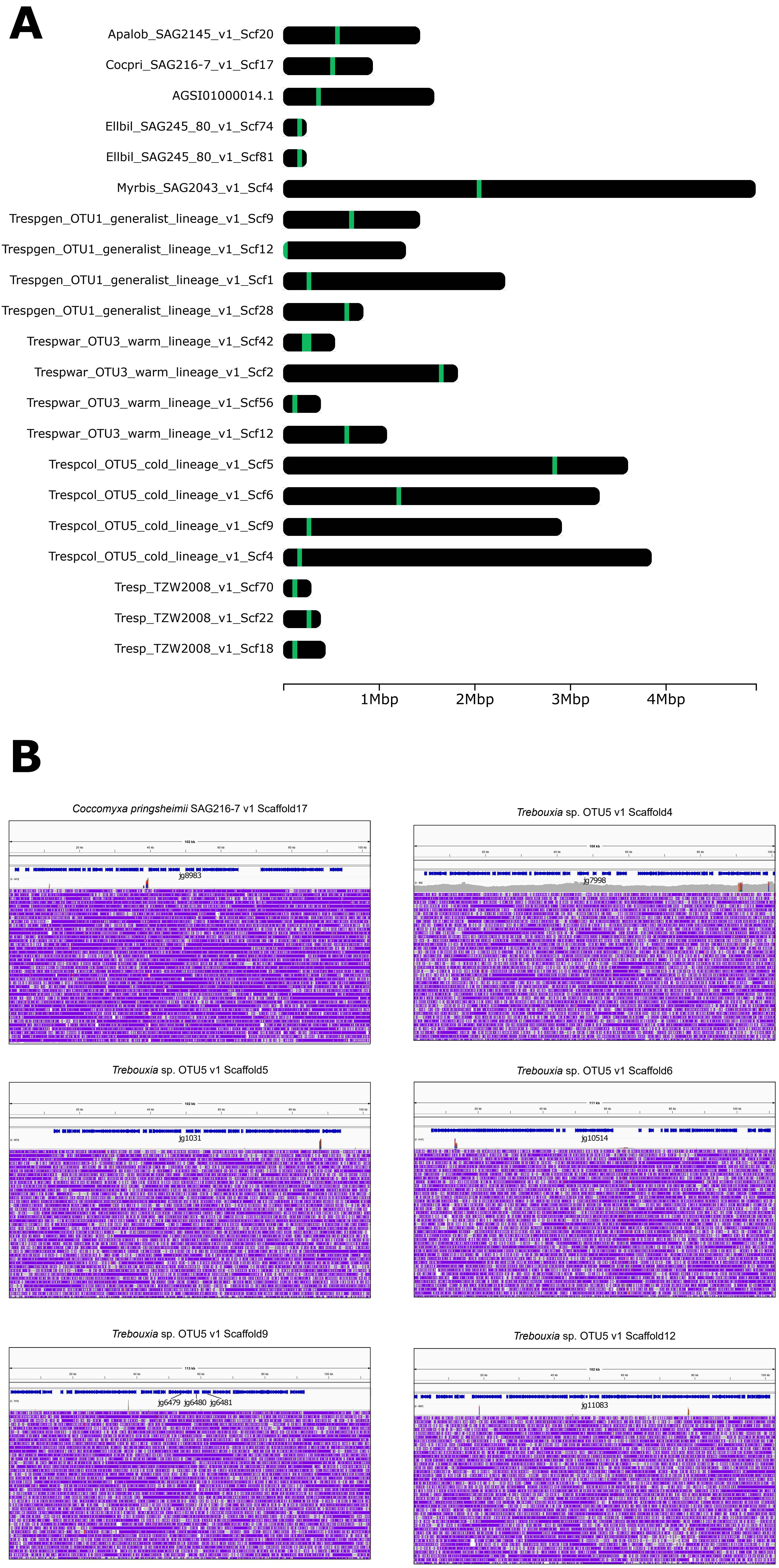


**Supplementary Fig. 9: Horizontal gene transfer demonstration for the glutathione S-transferase.** Read mapping and scaffold anchoring of the GST for *Coccomyxa pringhseimii* SAG216-7 and *Trebouxia sp* OTU1
